# A comparative analysis of *Caenorhabditis* and *Drosophila* transcriptional changes in response to pathogen infection

**DOI:** 10.1101/2020.03.04.977595

**Authors:** Robert L. Unckless, Patrick A. Lansdon, Brian D. Ackley

## Abstract

*Drosophila melanogaster* and *Caenorhabditis elegans* are well-used invertebrate models for studying the innate immune system. The organisms are susceptible to bacterial pathogens that include *Pseudomonas* species, *(entomophilia – Drosophila) or (aeruginosa – Caenorhabditis), E. faecalis* and *P. rettgeri*, which are or are related to human pathogens. Further, the consequences of exposure to these pathogens, in terms of organismal survival, are roughly equivalent when compared. That is, worms and flies are more susceptible to infection by *Pseudomonas* than *E. faecalis*, whereas organismal survival on *E. faecalis* and *P. rettgeri* are roughly the same in both. To better understand how these organisms are coordinating their responses to these bacterial pathogens we examined transcriptomes in infected animals. We grouped our analysis based on protein orthology. Of the 3611 pairs analyzed, we found genes whose responses were conserved across the different species at a higher than expected rate for two of the three pathogens. Interestingly within the animals, genes with 1:1 orthologs between species behaved differently. Such genes were more likely to be expressed in *D. melanogaster*, and less likely to be expressed in *C. elegans*. From this analysis we found that the gene nucleobindin (*nucb-1/NUCB1* in *C. elegans* and *D. melanogaster*, respectively) was upregulated in both species in response to Gram negative bacteria. We used RNAi to knock down *nucb-1* and found the treated animals were more susceptible to infection by the Gram negative pathogen *P. rettgeri* than controls. These results provide insight into some of the conserved mechanisms of pathogen defense, but also suggest that these divergent organisms have evolved specific means to orchestrate the defense against pathogens.

**Article Summary:** We analyzed transcriptomic data from *C. elegans* and *D. melanogaster* to compare the expression of orthologous pairs of genes in response to bacterial pathogens. Our results indicated that only a handful of genes that are orthologous between species are differentially expressed in response to pathogens, but that the pattern of expression was different when comparing one-to-one orthologs versus those that are restricted to one of the two organisms. These results suggest that, although broad patterns of susceptibility to bacterial pathogens are conserved, the regulatory framework by which the organisms fight pathogens is less well conserved. Further our results suggest a more complete analysis of the evolutionary changes in organismal responses to pathogens is required.

## Introduction

The innate immune system is ancient and evolutionarily conserved across the animal kingdom. By contrast the adaptive immune system evolved in bony fish about 500 million years ago (Boehm and Swann 2014). The function of the innate immune system is to protect organisms against pathogens such as bacteria, fungi and viruses. The broad repertoire and rapid evolution of pathogens appears to have had a compensatory effect in the adaptation of the innate immune system in animals. From this, there has been a general observation that immune genes are often amongst the fastest evolving genes in genomes (Nielsen *et al.* 2005; Sackton *et al.* 2007; Pujol *et al.* 2008; Downing *et al.* 2009; Obbard *et al.* 2009; Viljakainen *et al.* 2009; Shultz and Sackton 2019).

However, more recent work has suggested that evolution in the innate immune system is more nuanced than that. The innate immune system can be roughly divided into pathways that respond to bacterial pathogens (with differences in Gram-positive vs. Gram-negative), fungal pathogens and viruses. When examined within the different pathways, there is evidence for different signatures of adaptation between *Drosophila* species. Rapid evolution in immune genes seems to be specifically to the viruses they face (Duxbury *et al.* 2019; Hill *et al.* 2019) or in response to other ecological pressures (Hanson *et al.* 2019). The evolution of immune genes in *Caenorhabditis* species is less studied and recurrent positive selection appears to be much less pronounced (Pujol *et al.* 2008; Schulenburg *et al.* 2008; Dierking *et al.* 2016).

*Caenorhabditis elegans* (roundworms) and *Drosophila melanogaster* (fruit flies) have been important models for identifying fundamental mechanisms of how organisms fight infection(Buchon *et al.* 2014; Ermolaeva and Schumacher 2014). Work from these systems have uncovered genes and pathways, many that are conserved in humans, that facilitate response to infection. Primarily, these include the Toll and Imd pathways (flies) and insulin/FOXO and MAP Kinase pathways (worms). However, the Toll and Imd pathways, while well-conserved from flies to humans, are largely absent in nematodes (Schulenburg *et al.* 2008). In flies the insulin/FOXO pathway is important, but less critical to the overall response.

However, reviewing the work done suggests that there has been more limited comparison of information obtained from these two systems. Interestingly, where it has been measured, pathogens exhibit roughly similar patterns of virulence in these organisms. For example, *Pseudomonas* species are more virulent than *Enterococcus* or *Providencia*, whereas *Enterococcus* and *Providencia* are roughly equivalent (Sim and Hibberd 2016; Martin *et al.* 2017; Archer and Phillips 2018; Troha *et al.* 2018; Vasquez-Rifo *et al.* 2019), B.D. Ackley, Personal Observation). This suggests that, even if the animals have invested different genetic resources in fighting these pathogens, the outcomes have not significantly improved. We find that genes differentially induced after exposure to Gram-positive and Gram-negative microbes are more likely to be lineage restricted in *Drosophila* but one-to-one orthologs (with *Drosophila*) in *Caenorhabditis.* Furthermore, there is weak, but significant correlation between fold change (between Gram-negative to Gram-positive exposure) between flies and worms. One particular gene, *Nucb-1*, was significantly biased toward expression after Gram-negative infection, and appears to play a role specifically in fighting Gram-negative infections.

## Methods

### Data acquisition: Drosophila melanogaster

All *D. melanogaster* sequencing data was published previously (Troha *et al.* 2018). We downloaded data from the sequence read archive (SRA) for 12 hours post infection with *Enterococcus faecalis*, *Escherichia coli*, *Providencia rettgeri* and *Pseudomonas entomophila.* Details for all sequencing data sets is presented in Table S1.

### Data acquisition: Caenorhabditis elegans

*C. elegans* (N2, var. Bristol) were reared on *Escherichia coli* (OP50) on nematode growth media (NGM) plates using standard conditions (Brenner 1974). For pathogen exposure, NGM plates were seeded with 250 μL of bacteria, *Pseudomonas aeruginosa* (PA14)*, Enterococcus faecalis* (OG1RF), *Providencia rettgeri* (D. mel isolate) or OP50 (control), and incubated overnight at 37°C. Approximately 2,000 L4 stage worms were transferred to prepared NGM plates and incubated at 20°C for 24 hours. After 24 hours, worms were washed with M9 buffer three times to remove any residual bacteria. The worms were resuspended in 100 μL of M9 buffer and mechanically disrupted in liquid nitrogen using a ceramic mortar and pestle. Frozen tissue was transferred to a microcentrifuge tube and 1mL TRIZOL reagent was added. The tubes were flash frozen in liquid nitrogen and stored at −80°C. Three biological replicates were collected for each worm and pathogen strain.

Frozen worm tissue was thawed, vortexed for 5 seconds and incubated at room temperature for 5 minutes. After the addition of 470 μL chloroform (mixed by inversion and phase separated for 2 minutes at room temperature), the samples were centrifuged at 15,000 RPM at 4°C for 15 minutes. The upper aqueous phase containing RNA (approximately 600 μL) was transferred to a new RNase-free Eppendorf tube. Total RNA extraction was carried out using the Monarch RNA Cleanup Kit (New England Biolabs) according to the manufacturer’s instructions. Total RNA was quantified using the Qubit (Thermofisher Scientific) and RNA quality and integrity was assessed using the Agilent Tapestation 2200 (Agilent Technologies). Sequence libraries were prepared using the TruSeq Stranded mRNA Library Prep Kit (Illumina) and sequenced using single-end 1×75 bp sequencing on the Illumina NextSeq 550 platform.

#### Gene expression quantification

Raw reads were mapped to indexed transcriptomes using Kallisto version 0.46.0 (Bray *et al.* 2016) then total counts were rounded to the nearest integer and imported into R version 5.1.3 (R Development Core Team 2015) for analysis. We used all mRNA transcripts from *D. melanogaster* FlyBase release r6.30 (Thurmond *et al.* 2019) and the longest transcripts from *C. elegans* WormBase release WBcel235 (Harris *et al.* 2020) as references. We assessed differential expression of transcripts using DESeq2 (Love *et al.* 2014).

#### Creation of ortholog lists

We obtained a list of orthologs shared between *D. melanogaster* and *C. elegans* from InParanoid8 (Sonnhammer and Ostlund 2015). Briefly, we navigated to the “Browse the Database” page, selected *Drosophila melanogaster* and *Caenorhabditis elegans* as the two species and employed a custom script to both parse the data and distinguish between one-to-one orthologs and one-to-many orthologs.

#### Experimental infections in Drosophila

To determine the role of *NUCB1* in immune defense, we crossed a fat body / hemocyte driver (C564 – Bloomington Drosophila Stock Center #6982) to either a *NUCB1* RNAi construct (Bloomington Drosophila Stock Center #44019) or empty cassette (Bloomington Drosophila Stock Center #36304). All flies were maintained on a standard molasses diet at 23°C with a 24-hour light cycle (12 hours light followed by 12 hours dark). Adults were transferred to new food vials after eclosion and males were infected at three to six days old. All infections took place in the morning (between 1 and 2 hours after “daylight”). Infections were performed by septic injury using a minutien pin dipped in either sterile Luria Bertani media (control), *P. rettgeri* (diluted to OD_600_ = 1.00 ± 0.02), or *E. faecalis* (diluted to OD_600_ = 1.5 ± 0.02). For survival assays, the number of flies alive was noted every morning for seven days. For bacterial load, we homogenized groups of three flies in 1X PBS, diluted 1:20, then plated the dilute homogenate on LB agar using the WASP Touch spiral plater (Don Whitley Scientific, West Yorkshire, UK). These plates were incubated at 37°C for about 18 hours then counted using a Flash & Go Plate Counter (Neu-tec Group Inc., Farmingdale, NY, USA).

#### Experimental infection in C. elegans

Wild-type (N2) animals were reared on *nucb-1* RNAi-expressing bacteria (Source Bioscience, Nottingham, UK). *nucb-1* RNAi bacteria were grown overnight at 37°C in 1.5 mL LB plus ampicillin, tetracycline and 1 mM IPTG with shaking. 0.2 ml of the overnight cultures were aliquoted onto NGM plates containing carbenicillin, tetracycline, and 1mM IPTG and grown overnight at 37°C. Healthy L4 animals were transferred to the *nucb-1* RNAi plates or OP50 (*E. coli*) control plates and allowed to produce offspring. Offspring were transferred to freshly prepared plates and reared for 2 days. 20-30 L4 animals from the RNAi or control plates were transferred to NGM plates seeded with *E. coli, E. faecalis* or *P. rettgeri* and then maintained at 20°C for the remainder of the experiment. Every 24 hours living animals were transferred to NGM plates with the same bacterial species and the number of living and dead animals was counted. Animals were transferred to fresh plates daily, until all animals were dead. RNAi was done in triplicate.

## Results

### One-to-one orthologs are less likely to be differentially expressed than other genes in worms but not flies

We measured gene expression after four different treatments in both *Drosophila* and *Caenorhabditis.* Principal component analysis of gene expression reveals that in both flies and worms, the sole Gram-positive pathogen, a moderately virulent strain of *Enterococcus faecalis*, clustered distinctly from other samples (Figure S1). In *Drosophila*, the two pathogenic Gram-negative microbes (*Pseudomonas entomophila* and *Providencia rettgeri*) clustered together while the non-pathogenic *Escherichia coli*) was somewhat intermediate between *E. faecalis* and the pathogenic Gram-negative pathogens. In *Caenorhabditis, E. coli* is the food source so we would expect very little immune response. Strangely, however, *E. coli* clusters with the relatively virulent *P. rettgeri* while the very virulent *Pseudomonas aeruginosa* is quite different. Thus, it appears that both broad bacterial taxonomy and virulence drive expression similarities in both flies and worms, but are not the only factors involved.

Since two of the most highly immune-induced pathways in *Drosophila* (*Imd* and *Toll*) do not exist in *Caenorhabditis*, we hypothesized that immune induction would tend to involve lineage-specific genes and not conserved orthologs. We therefore assigned all genes in each taxa as lineage restricted, one-to-many orthologs or one-to-one orthologs (with orthology considered compared to the other taxa: flies to worms and worms to flies). This resulted in 3543/3580 one-to-one orthologs, 3428/3298 one-to-many orthologs and 10854/9102 lineage restricted genes in *Drosophila* and *Caenorhabditis*, respectively. In flies, there were 79, 331 and 248 differentially expressed genes comparing *E. faecalis* to *E. coli, P. aeruginosa* and *P. rettgeri*, respectively (FDR=0.05). In worms, there were 5867, 6375 and 5868 differentially expressed genes comparing *E. faecalis* to *E. coli*, *P. aeruginosa* and *P. rettgeri*, respectively (FDR=0.05). In *Drosophila*, lineage restricted genes were most likely to be differentially expressed, followed by one-to-many orthologs, then one-to-one orthologs when comparing infection with *E. faecalis* to any of the Gram-negative bacteria (Figure 1A). In all cases, orthology category was significantly associated with the likelihood of being differentially expressed, however, Tukey honestly significant difference tests were significant for only some pairwise comparisons (Supplemental Table 2). The pattern for *Caenorhabditis*, however, is quite different: lineage restricted genes are least likely to be differentially expressed, followed by one-to-many orthologs and one-to-one orthologs are most differentially expressed (Figure 1B, Supplemental Table 2).

**Figure 1:**
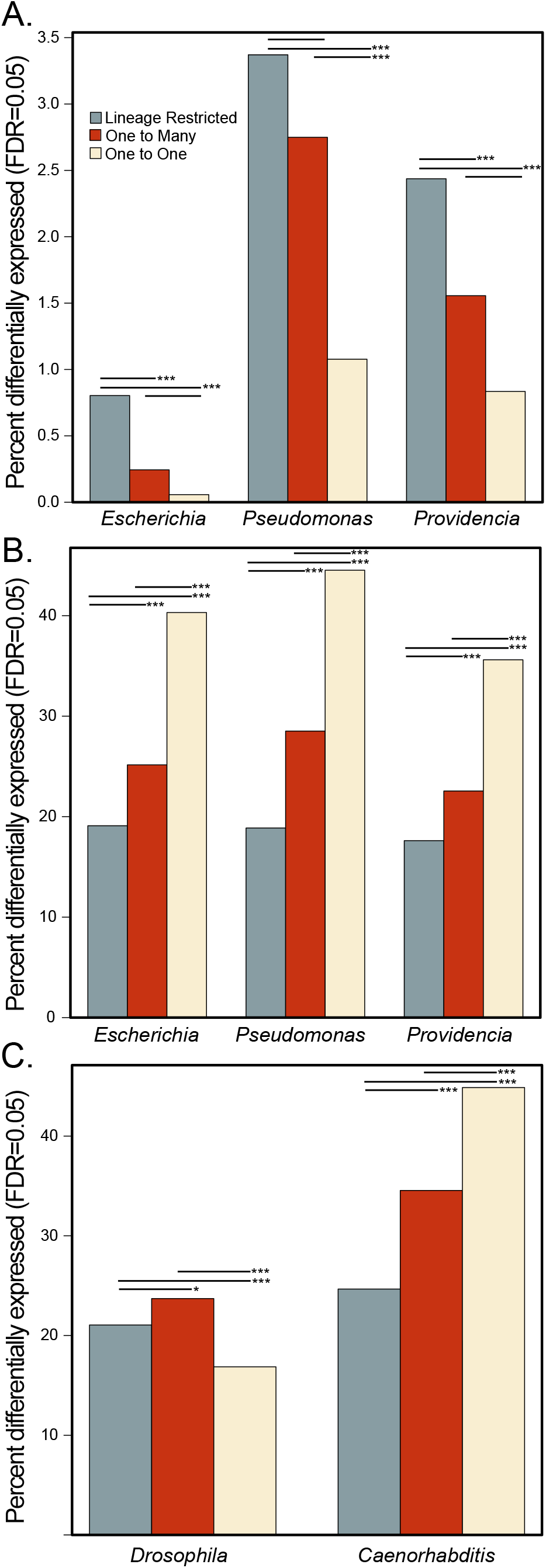
Differentially expressed genes are associated with orthology status in both *Drosophila* and *Caenorhabditis* but in opposite directions. The proportion of differentially expressed genes (FDR=0.05) in A) *Drosophila* and B) *Caenorhabditis* exposed to *E. faecalis* vs. *Escherichia*, *Pseudomonas* and *Providencia* (see supplemental table S?? for strain names, and c) both *Drosophila* and *Caenorhabditis* control compared to *Pseudomonas*. *P<0.05, **P<0.001, ***P<0.0001.

**Figure 2:**
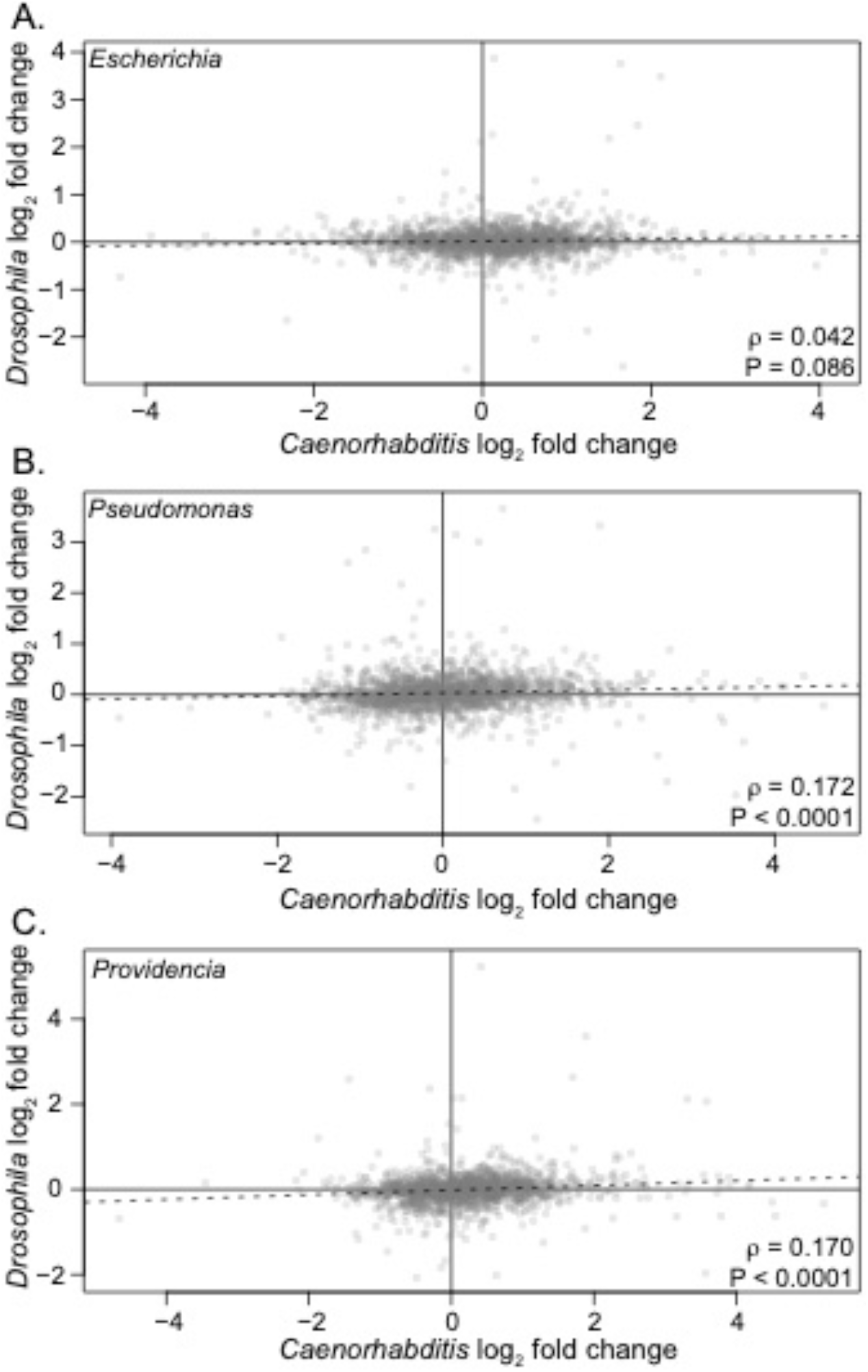
Correlations between *Drosophila* and *Caenorhabditis* log_2_ fold change between *E. faecalis* and A) *E. coli*, B) *Pseudomonas aeruginosa or entomophila* or C) *P. rettgeri*. Spearman’s rho value and P-values are presented in the lower right hand corner of each plot.

### Expression differences between individuals exposed to Gram+ and Gram− pathogens are weakly correlated in one-to-one orthologs in divergent hosts

When considering only one-to-one orthologs, the log_2_ fold change after exposure to *E. faecalis* vs. all three Gram-negative pathogens was positively correlated, but weakly. The correlation for *E. coli* was weakest (*ρ* = 0.042, P = 0.086), with both *Pseudomonas* (*ρ* = 0.172, P < 0.0001) and *P. rettgeri* (*ρ* = 0.170, P < 0.0001) similar to each other, but only moderately correlated. This suggests that while there is some degree of conservation of the overall difference in expression response to Gram-positive vs. Gram-negative pathogens, the pattern is relatively weak. A handful of genes were significantly differentially expressed (in the same direction) in both flies and worms when comparing exposure to *E. faecalis* to *Pseudomonas* or *Providencia* (Table 1).

**Table 1:** Genes significantly differentially expressed in both *Drosophila* and *Caenorhabditis*.

| Pathogen | Gene | *Drosophila* |  |  | Gene | *Caenorhabditis* |  |  |
| --- | --- | --- | --- | --- | --- | --- | --- | --- |
|  |  | Base | Log<sub>2</sub> | P<sub>adj</sub> |  | Base | Log<sub>2</sub> | P<sub>adj</sub> |
|  |  | Count | FC |  |  | Count | FC |  |
| *Pseudomonas* | *Pu* | 888.6 | 2.60 | 4.28E-30 | *cat-4* | 82.1 | -1.14 | 8.75E-05 |
|  | *aay* | 1028.1 | 1.50 | 1.18E-05 | *Y62E10A.13* | 402 | -0.35 | 1.88E-02 |
|  | *Dgp-1* | 153.0 | 1.09 | 1.26E-03 | *cgp-1* | 1687 | -0.6 | 2.11E-10 |
|  | **CG14210** | **17.8** | **-1.81** | **3.85E-02** | **Y40B1B.7** | **810.3** | **-0.39** | **1.18E-03** |
|  | *CG9643* | 66.0 | 0.98 | 2.70E-02 | *F29B9.1* | 410.6 | -1.15 | 1.84E-15 |
|  | *P32* | 713.1 | 2.84 | 1.28E-24 | *cri-3* | 1702.3 | -0.93 | 1.47E-15 |
|  | *CG10103* | 149.4 | 0.89 | 8.29E-04 | *istr-1* | 1181.6 | -0.29 | 4.44E-03 |
|  | **NUCB1** | **957.5** | **0.89** | **4.02E-07** | **nucb-1** | **132.5** | **1.47** | **3.71E-08** |
|  | **Galt** | **302.5** | **-0.91** | **3.85E-02** | **ZK1058.3** | **331.5** | **-0.97** | **4.79E-10** |
|  | **Esyt2** | **158.5** | **0.81** | **5.69E-04** | **esyt-2** | **149.2** | **1** | **2.20E-05** |
| *Providencia* | **ade2** | **149.9** | **-1.03** | **2.09E-04** | **pfas-1** | **61.4** | **-1.34** | **1.28E-03** |
|  | *Pu* | 888.6 | 2.59 | 5.36E-30 | *cat-4* | 82.1 | -1.44 | 8.37E-06 |
|  | *P32* | 713.1 | 2.36 | 1.28E-16 | *cri-3* | 1702.3 | -0.31 | 2.45E-02 |
|  | **CG6218** | **360.0** | **0.51** | **4.47E-02** | **W06B4.2** | **372.3** | **0.39** | **2.15E-02** |

#### Genes differentially expressed comparing infection with Pseudomonas to E. faecalis

*CG14210*/*Y40B1B.7* was Gram-positive biased in both animals but is poorly characterized and is an ortholog of the human *CCDC86* gene (coiled-coil domain containing 86) which localizes to the nucleolus (Thurmond *et al.* 2019; Harris *et al.* 2020). *NUCB1*/*nucb-1* was Gram-negative biased in both animals and is a nucleobindin protein involved in defense against Gram-negative bacteria in *Drosophila* (Berkey et al. 2009) and is expressed in the intestines of *Caenorhabditis* (Harris *et al.* 2020). *Galt*/*ZK1058.3* was Gram-positive biased in both animals is a galactose-1-phosphate uridylyltransferase associated with galactose homeostasis in *Drosophila* and without phenotypic data in *Caenorhabditis* (Thurmond et al. 2019). In addition, *Galt* was upregulated in *Drosophila* after infection with *Octosporea* (a microsporidian that normally infects *Daphnia*) (Roxstrom-Lindquist *et al.* 2004). *Esyt2*/*esyt-2* was Gram-negative biased and is a synaptotagmin-like protein involved in lipid transport, among other things (Thurmond *et al.* 2019; Harris *et al.* 2020).

#### Genes differentially expressed comparing infection with P. rettgeri to E. faecalis

*ade2*/*pfas-1* was significantly Gram-positive biased after exposure and is a phosphoribosylformylglycinamidine synthase and is involved in purine synthesis and is expressed in the fat body in *Drosophila* and the intestines in *Caenorhabditis* (Thurmond *et al.* 2019; Harris *et al.* 2020). *CG6218*/*W06B4.2* was gram-negative biased in both animals and is a N-acetylglucosamine kinase involved in carbohydrate phosphorylation and influences dauer lifespan in *Caenorhabditis* (Xie and Roy 2012).

### GO Analysis reveals different pathways influenced by infection

Several Gene Ontology categories were enriched when comparing exposure to the Gram-positive *E. faecalis* to the three Gram-negative microbes (Supplemental Table 3). Focusing on molecular function, the many categories enriched for one pathogen were also enriched for other categories in both *Drosophila* (Figure 3A) and *Caenorhabditis* (Figure 3B), with the important caveat that in all cases the three different Gram-negative microbe treatments are being compared to the same Gram-positive *E. faecalis* treatment. Overlaps are significantly more common than expected by chance in both species in all pairwise comparisons (Supplemental Table 4). However, the only molecular function Gene Ontology category shared between *Drosophila* and *Caenorhabditis* after exposure to any of the microbes was “GO:0001055 - RNA Polymerase II Activity”.

**Figure 3.**
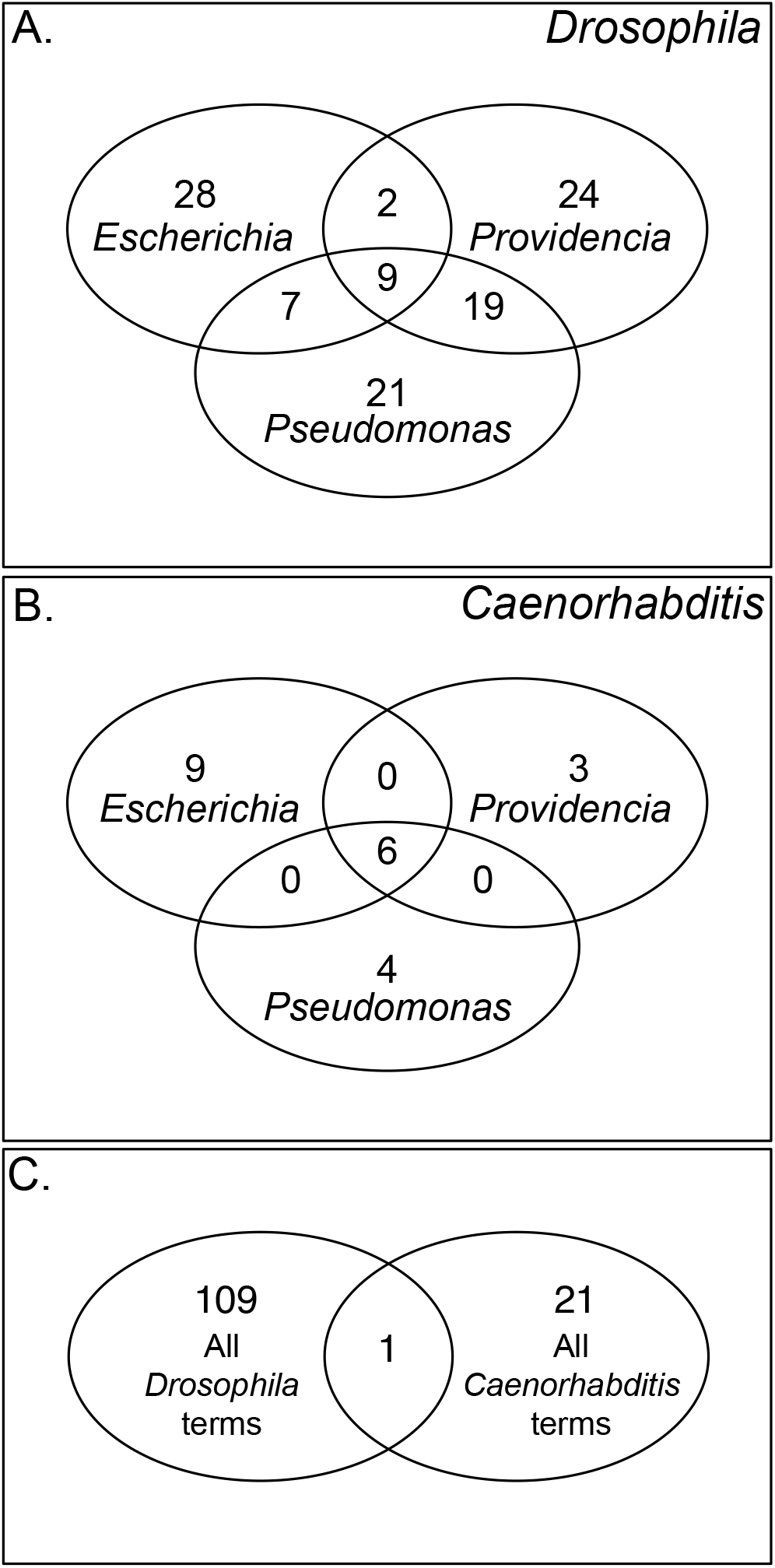
Gene Ontology term enrichment overlap between comparing Gram-positive *E. faecalis* to the three Gram-negative microbes. A) *Drosophila*, B) *Caenorhabditis* and C) all significant *Drosophila* Molecular Function terms to all significant *Caenorhabditis* Molecular Function terms.

### Oral infection in Drosophila is largely consistent/inconsistent with previous results

Since the above analysis compares systemic infection in *Drosophila* to oral infection in *Caenorhabditis*, we also compared *Drosophila* gene expression 16 hours after oral infection with *P. entomophila* (Bou Sleiman et al. 2020). However, in this analysis guts, not whole animals were used as a source of RNA. Also, in this case, we compared control expression (standard diet) to expression 16 hours after exposure to *P. entomophila* in *Drosophila*. The best contrast in *Caenorhabditis* was comparing exposure to *E. coli* (which is the standard worm diet) to exposure to *P. aeruginosa.* So in this section, for both taxa, we are comparing standard diet to a highly virulent oral *Pseudomonas* infection. Like *Drosophila* systemic infection, lineage restricted genes (21.1%) were significantly more likely to be differentially expressed than one-to-one orthologs (16.9%), but one-to-many orthologs were actually the most likely class to be differentially expressed (23.7%, Table S2). In this new comparison (*Pseudomonas* vs. *Escherichia*) in *Caenorhabditis*, the patterns of differential expression and orthology look very much like *Pseudomonas* vs. *E. faecalis*, suggesting that the patterns of differential expression might be driven by the virulence of *Pseudomonas*.

The log_2_ fold change of one-to-one orthologs after oral infection was negatively correlated between *Drosophila* and *Caenorhabditis* (*rho*=−0.087, P<0.0001), suggesting that perhaps the whole-animal vs. gut-only approach led to this negative correlation. A handful of genes were significantly induced in both animals and include *cv-2/T01D3.6*, *Jra/jun-1*, *vri/atf-2*, *alpha-Man-IIb/aman-3*, *CG10681/kxd-1*, *spds/spds-1*, *CG10184/R102.4*, and *CG32549/Y71H10B.1* (fly gene symbol/worm gene symbol). *Jra/jun-1* is of particular interest because it is a negative regulator of NF-KappaB mediated activation of antimicrobial peptides in *Drosophila* and mutant heterozygotes had reduced survival after infection with generally avirulent *E. coli* (Kim et al. 2007). Interestingly, *jun-1* mutants in *C. elegans* had increased survival after infection with *P. aeruginosa* (Kao et al. 2011). *alpha-Man-IIb/aman-3* has alpha-mannosidase activity and is associated with the encapsulation response in *Drosophila* (Mortimer et al. 2012).

No gene ontology terms were significantly overrepresented in both *Drosophila* and *Caenorhabditis* (Supplementary Table 4).

### Experimental infections reveal a Gram-negative specific role in immune defense for Nucb-1

The *nucb-1 / NUCB1* Orthologs were significantly Gram-negative biased in both flies and worms. This is interesting since the Toll and IMD pathways that regulate defense against Gram-positive and Gram-negative microbes in *D. melanogaster* are missing in *C. elegans.* We therefore tested whether knocking down *nucb-1* would show an immune phenotype after challenge with *P. rettgeri* or *E. faecalis*. First, there was a significant interaction between treatment and genotype on log-transformed total bacteria per fly (Figure 4A, Supplementary Table 5, P=0.031) 24 hours after infection. The mean *E. faecalis* load for males with *NUCB1* knocked down was 8% of controls (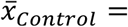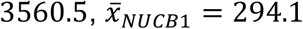). However, the mean *P. rettgeri* load for males with *NUCB1* knocked down was more than 400% of controls (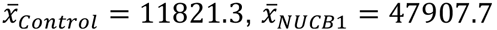). We found a similar interaction effect (P=0.012, Supplementary Table 6) when analyzing survival after infection (Figure 4B). *NUCB1* knockdown males survived slightly better than controls after infection with *E. faecalis* (77.8% controls vs. 83.3% *NUCB1* survived five days post infection) but had lower survival after infection with *P. rettgeri* (83.3% controls vs. 66.7% *NUCB1* survived four days post infection). These results are qualitatively similar when analyzing using a different day after infection or a Cox Proportional Hazard framework (data not shown). In *C. elegans, nucb-1* knockdown individuals survived *E. faecalis* exposure better than untreated individuals (78.8% knockdown vs. 32.2% untreated, Figure 4C). In contrast, *Nucb-1* knockdown individuals survived *P. rettgeri* exposure worse than untreated individuals (38.8% knockdown vs. 66.7% untreated). Again, the interaction between treatment (microbe exposure) and knockdown was significant (P<0.0001, Supplemental Table 7).

**Figure 4.**
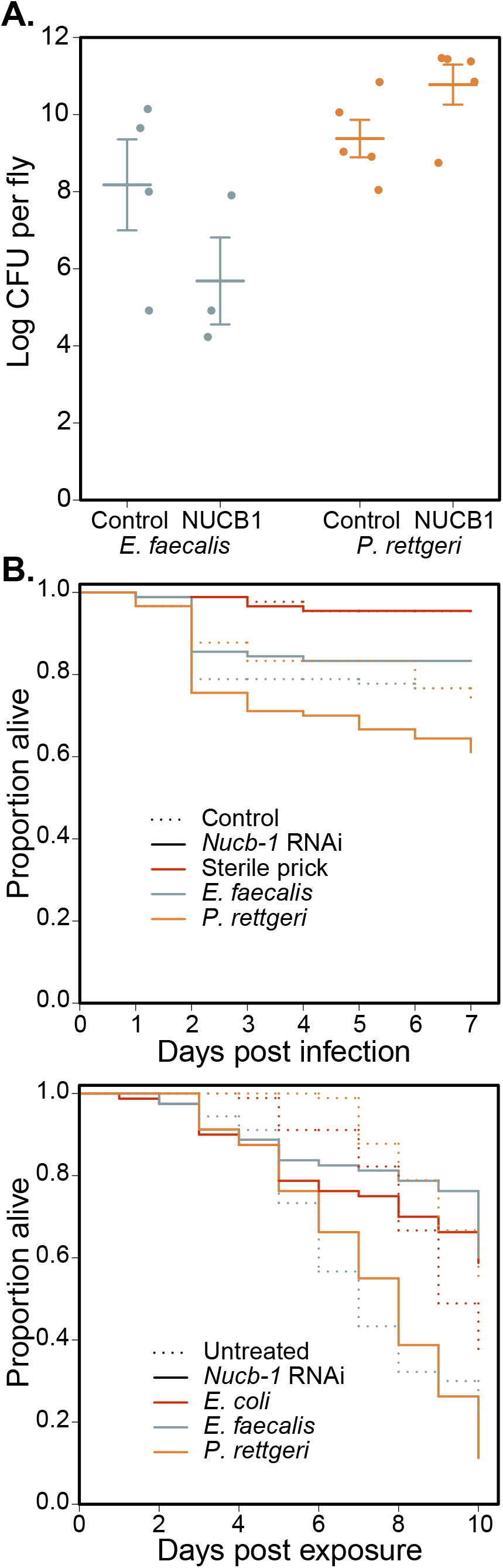
*Nucb-1* protects specifically against Gram-negative pathogens in flies and worms. A) Bacterial load presented as natural log colony forming units (CFU) per individual fly 24 hours after infection with *E. faecalis* or *P. rettgeri* in control (fat body driver with empty cassette) or *Nucb-1* knockdown (fat body driver with *Nucb-1* RNAi construct). Thick bars represent means and error bars represent plus or minus one standard error of the mean. B) Survival after sterile prick or infection with *E. faecalis* or *P. rettgeri* in control (fat body driver with empty cassette) or *Nucb-1* knockdown (fat body driver with *Nucb-1* RNAi construct). n=90 for each sample. C. Survival after exposure to *E. coli* (standard conditions), *E. faecalis* or *P. rettgeri* in untreated or *nucb-1* knockdown (worms fed *E. coli* expressing *nucb-1* RNAi construct). n=80-90 for each sample.

## Discussion

*D. melanogaster* and *C. elegans* are arguably the two most well-studied invertebrate model systems. Interestingly, their ecologies are overlapping and *C. elegans* can even be phoretic on *D. melanogaster* (Kiontke and Sudhaus 2006). Because of this shared ecology, we would expect that the microbes each encounter in nature are also intersecting and therefore that immune defense against these microbes would be under similar selective pressures in the two models. However, innate immunity in flies and worms appears quite different. *Caenorhabditis* lacks the canonical NF-KappaB pathways (Toll and Imd in *Drosophila*). We performed a first step in a systematic comparison how the two species defend themselves against pathogenic microbes using transcriptomic data and comparing exposure to Gram-positive *E. faecalis* to exposure to three different Gram-negative microbes. We find that lineage restricted genes are more likely to be differentially expressed between exposure types in *Drosophila*, but one-to-one orthologs are more likely to be differentially expressed in *Caenorhabditis.* Further, among conserved, one-to-one orthologs, there was weak but significant correlation between *Drosophila* and *Caenorhabditis* in the log_2_ fold change between exposure to *E. faecalis* and all three Gram-negative microbes. However, we found very little evidence that similar gene ontologies are enriched when comparing *E. faecalis* infection to the Gram-negatives. These results suggest that while flies and worms share similar ecologies and are therefore likely exposed to similar microbes, they seem to fight infection in very distinct ways.

One obvious caveat is that our comparison is, necessarily, not perfect. Most of the *Drosophila* data utilized were from systemic infection involving pinprick with a septic needle whereas *Caenorhabditis* were fed microbes leading to primarily gut infection. Even when using *Drosophila* gut infection data, we are comparing whole organism (worm) to dissected guts (flies). Finally, the timeframe for infection dynamics could be very different in the two hosts and this difference could be exacerbated by the different routes of infection. We feel, however, that the results are probably not a result of these differences in approach. In both flies and worms, infections are strongly induced by the timepoints analyzed and those time points are during the critical period when infection outcomes are decided (Duneau *et al.* 2017). Instead, we think that the differences are more likely due to fundamentally different pathways governing immune defense in the two hosts.

If host responses are fundamentally different, the specific role that a few key genes (i.e. nucleobindin) play in both species is particularly interesting. If *Nucb-1* is a Gram-negative specific immune protein in both worms and flies, but uses non-canonical pathways, it might suggest a new conserved immune pathways in these two disparate hosts. *NUCB1* was previously implicated in *Drosophila* in immune defense, but not in *Caenorhabditis* (Berkey *et al.* 2009). We found that in both organisms immune defense against the Gram-negative pathogen, *P. rettgeri* is significantly reduced when *Nucb-1* expression is knocked down. Moreover, immune defense against *E. faecalis* seems to improve when *Nucb-1* expression is knocked down. This motivates a deeper study of the roll of *Nucb-1* in immune defense in both hosts: mechanisms of specific defense, regulation and upstream and downstream interactors.

Overall, our results represent an initial attempt to compare immune defense pathways between *C. elegans* and *D. melanogaster.* This work has identified genes that may be good candidates for underlying the similarities of immune defense in organisms that can share an ecological niche. Moreover, the pathogens we have used here represent bacteria that are, or are related to, human health hazards. Identifying mechanisms that are conserved between flies and worms may help us to better identify mechanisms that function in human immunity. In the long term, it may be possible to use the induction of these genes in people to evaluate infection and whether the host is responding in the most efficient manner.

## Supplemental Tables

**Supplemental Table 1:** SRA accessions and other properties of sequencing runs for this study

**Supplemental Table 2: Logistic regression and Tukey HSD for orthology categories.** LR: logistic regression, RD: residual deviance, d.f.: degrees of freedom, P(1:many – LinRes): Tukey HSD P-value for 1 vs. many orthologs compared to lineage restricted orthologs, etc.

**Supplemental Table 3:** List of all significantly enriched gene ontology (GO) terms

**Supplemental Table 4.** Significant overlap in enrichment of gene ontology (GO) terms

**Supplemental Table 5.** *Drosophila* bacterial load is influenced by *NUCB1*. Log CFU per fly analyzed by type 3 anova with genotype, treatment and their interaction as factors.

**Supplemental Table 6.** *Drosophila s*urvival after infection is influenced by *NUCB1*. Data analyzed using logistic regression of survival five days after infection.

**Supplemental Table 7.** *Caenorhabditis s*urvival after infection is influenced by *NUCB1*. Data analyzed using logistic regression of survival five days after infection.

## Supplemental Figures

**Figure S1:**
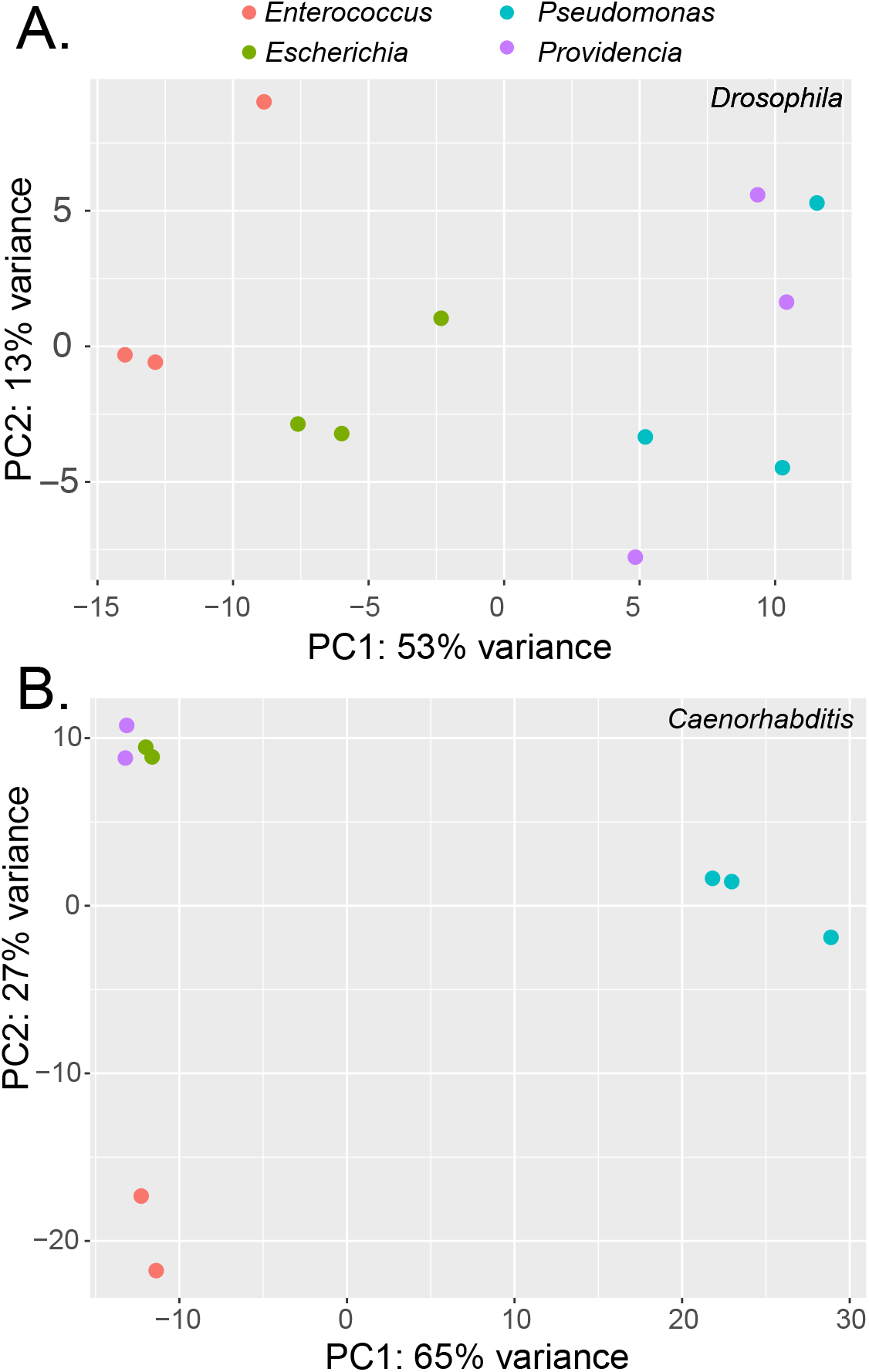
Principal component plots for A) *Drosophila* and B) *Caenorhabditis* samples. See supplemental table S1 and materials and methods for strain names.

